# Reverse engineering neuron type-specific and type-orthogonal splicing-regulatory networks using single-cell transcriptomes

**DOI:** 10.1101/2024.06.13.597128

**Authors:** Daniel F Moakley, Melissa Campbell, Miquel Anglada-Girotto, Huijuan Feng, Andrea Califano, Edmund Au, Chaolin Zhang

## Abstract

Cell type-specific alternative splicing (AS) enables differential gene isoform expression between diverse neuron types with distinct identities and functions. Current studies linking individual RNA-binding proteins (RBPs) to AS in a few neuron types underscore the need for holistic modeling. Here, we use network reverse engineering to derive a map of the neuron type-specific AS regulatory landscape from 133 mouse neocortical cell types defined by single-cell transcriptomes. This approach reliably inferred the regulons of 350 RBPs and their cell type-specific activities. Our analysis revealed driving factors delineating neuronal identities, among which we validated Elavl2 as a key RBP for MGE-specific splicing in GABAergic interneurons using an in vitro ESC differentiation system. We also identified a module of exons and candidate regulators specific for long- and short-projection neurons across multiple neuronal classes. This study provides a resource for elucidating splicing regulatory programs that drive neuronal molecular diversity, including those that do not align with gene expression-based classifications.

## Introduction

Normal brain function relies on a diverse repertoire of neurons, which together enable the emergent properties of neural circuits. Neuron types vary in many physiological characteristics, including their spatial distribution, the axonal inputs and targets, the neurotransmitters they release, and their electrophysiological properties^1^. Some of these phenotypic differences can be delineated by distinct developmental lineages. Alternatively, convergent or orthogonal mechanisms can dictate certain functional properties in specific neuronal subtypes across different lineages, suggesting a multifaceted array of factors determining neuronal identity and function. For example, glutamatergic excitatory neurons and GABAergic inhibitory interneurons are two principal neuronal classes with distinct morphological, functional and connective properties, reflecting their developmental origins in the pallium and subpallium^2–6^. On the other hand, neuron types with long-projecting axons can be found in both glutamatergic and GABAergic neuronal classes and these neurons have common biological needs associated with the upkeep of a long axon despite their difference in developmental lineages^7,8^.

At the molecular level, neuronal identity and function are specified by the selective activity of gene-regulatory programs. Much progress has been made in elucidating the tissue and cell type-specific patterns of regulation that drive the RNA transcription levels required for gene function. In particular, technological advancements in single-cell RNA sequencing (scRNA-seq) have enabled unbiased concurrent examination of all brain cell types, which has led to the identification of hundreds of “transcriptional cell types” in the mouse and human brain^8–11^ and the elucidation of the underlying transcriptional regulatory programs driven by DNA-binding transcription factors. However, the contributions of different steps of co- or post-transcriptional RNA processing to neuronal cell identity and function are less well studied.

Alternative splicing (AS) allows cells to customize gene products by selectively including or excluding exons from the mature RNA species, providing a major source of transcriptional diversity, especially in the brain^12–15^. In this process, RNA-binding proteins (RBPs) repress or activate the inclusion of target exons by binding to flanking *cis-*regulatory elements and interacting with the splicing machinery, thus altering the coding regions in the mRNA transcripts or fine-tuning mRNA stability, localization, and translation, as required by specific cellular contexts^16,17^. Although functions of most neuron type-specific AS events remain unknown, proper AS in neurons has been shown to be critical for neural development and physiology, providing modular roles for genes involved in, for example, neuronal migration, axonal assembly, cell adhesion signaling, and synaptic transmission^18–28^. Dysregulation of AS in the nervous system has been linked to a range of neurological disorders^17^, some of which are known to be caused by cell type-specific deficiencieş including SMA^29^, autism^21^, and schizophrenia^30^. A full understanding of how the molecular identities of neuron types are established, and how they may go awry in disease, will require elucidation of the AS regulatory landscape. Towards this goal, recent studies from several groups including us have used bulk or scRNA-seq data to catalog neuron type-specific AS^31–33^. Efforts have also been made to reveal RBP regulators that dictate neuron type-specific AS, but these studies have mostly focused on particular cell types and/or only a limited set of putative regulators^15,22,32–36^. The field still lacks a comprehensive and reliable map of the splicing-regulatory networks mediating AS differences amongst neuron types.

Here we present an end-to-end solution named Master Regulator analysis of Alternative Splicing (MR-AS) to reverse engineer splicing-regulatory networks using scRNA-seq data from diverse neuron types. MR-AS builds on the optimization of the ARACNe/VIPER framework we previously developed, which has been successfully employed to reverse engineer transcriptional regulatory networks in cancer and other cellular contexts using an information theoretic approach^37,38^, and identify master regulators by estimating protein activity from the aggregated behaviors of the target genes (i.e., regulons)^39,40^. Applying this pipeline to 133 adult mouse cortical cell types using AS and gene expression quantified in the same scRNA-seq data, we demonstrate that MR-AS can derive reliable splicing-regulatory networks and infer putative regulators that drive differential splicing between neuron types at different clades. We experimentally validated our computational predictions by focusing on the role of Elav-like 2 (Elavl2), a neuron-specific splicing factor, in directing differential splicing between medial ganglionic eminence (MGE)- and caudal ganglionic eminence (CGE)-lineage interneurons, the two primary subclasses of cortical inhibitory interneurons. We also identified a module of exons whose inclusion is specific to long-versus short-projecting neurons across a range of unrelated neuron types, as well as multiple RBPs, including Rbfox3, Khdrbs3, Qk and Elavl3, as potential driving factors. We show that reprogramming neuronal identity with altered axon length is accompanied by consistent AS switches of these module exons. Together, these results suggest the validity of the inferred splicing-regulatory networks and neuron type-specific RBP activities as a resource to elucidate the complex landscape of post-transcriptional gene regulation in the mammalian nervous system.

## Results

### Reverse engineering of splicing-regulatory networks and estimation of RBP activity

To investigate the rich diversity of AS regulation across neuron types, we leveraged the scRNA-seq data from 21,154 mouse neocortical cells generated by the Allen Institute for Brain Science^8^. This dataset used the SMART-seq protocol to achieve full-transcript read coverage, allowing simultaneous quantification of gene expression and splicing^15^. After experimenting with different strategies for imputing missing data and inferring splicing-regulatory networks, we decided to apply the ARACNe algorithm to cassette exon inclusion and RBP expression levels of 133 cell types using pseudobulk samples pooled from single cells to associate exons and RBPs using mutual information (Figure 1A,B and Figure S1; see Methods). After removing RBPs with less than 25 edges, the resulting network consisted of 174,274 edges between 350 RBPs and 8,336 cassette exons, with an average of 498 target exons per RBP, referred to as “RBP regulons” (Figure 1C and Table S1). For each putative target exon, the direction and likelihood of regulation by the RBP (mode of regulation; MOR) was determined by the direction and magnitude of RBP-exon correlation as described previously^39,40^. The number of targets per RBP followed approximately a power law distribution, indicating the scale-free property of the inferred network (Figure 1D). The inferred regulators of neuron type-specific exons between various clades of neuron types included a number of RBPs previously we have previously shown to play a role in neuron-specific AS regulation (Figure 1E and 1F; ref. ^15^).

**Figure 1.**
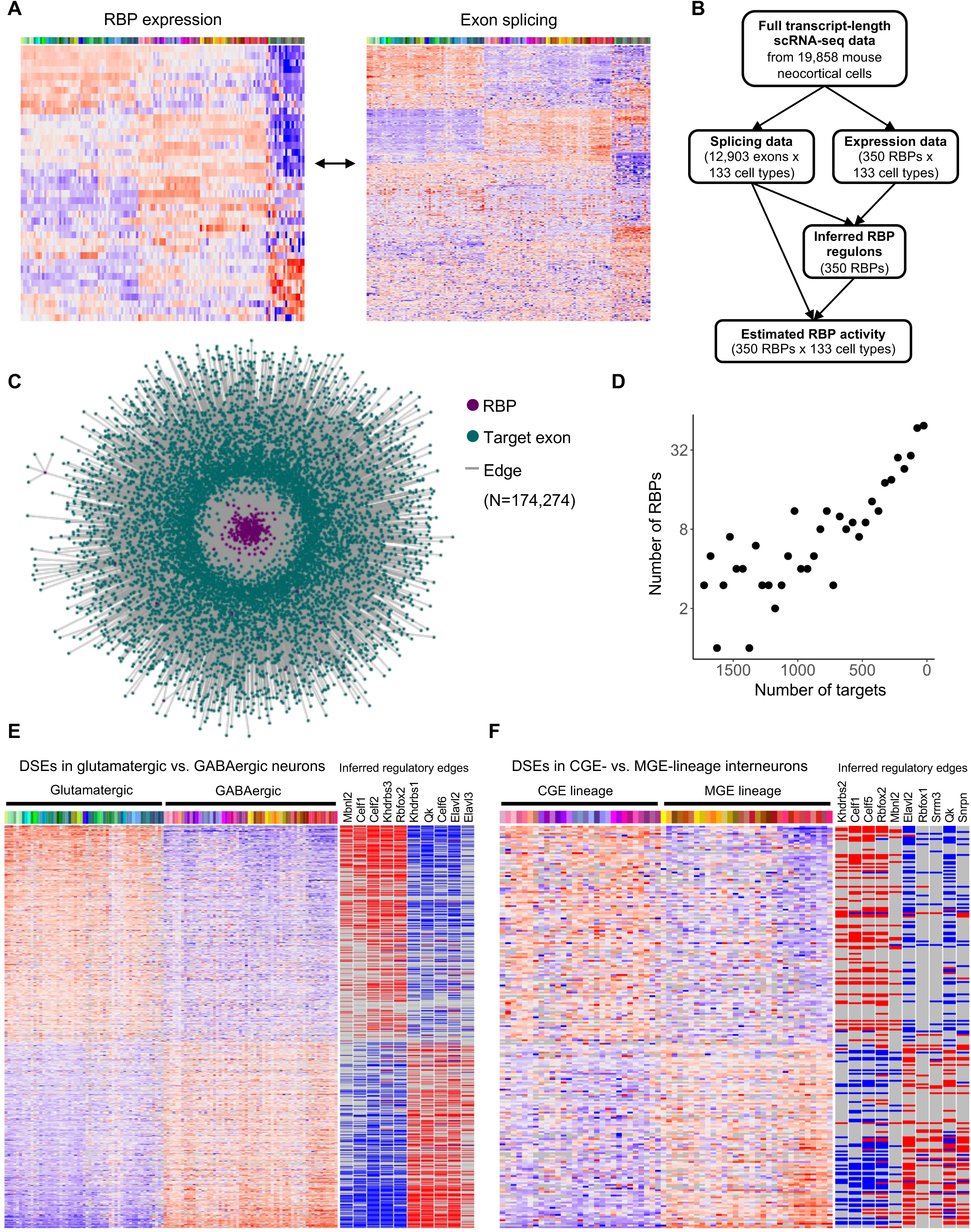
Reconstruction of splicing-regulatory networks using scRNA-seq data from adult mouse neocortical cell types. **A**, Adult mouse neocortical cell scRNA-seq data^8^ are pooled by cell types for the simultaneous quantification of RBP expression and cassette exon inclusion. **B**, Schematic of splicing regulation inference workflow. The ARACNe algorithm is applied to the cell-type-level RBP expression and exon inclusion data to predict the regulatory targets (regulon) of each RBP, which is then used to estimate neuron type-specific RBP activity using the VIPER algorithm. **C**, Visualization of the inferred splicing-regulatory network. **D**, Number of inferred targets per RBP. RBPs are grouped into 50 bins based on the regulon size. The inferred regulon size per RBP approximately follows a power-law distribution. **E**, Heatmap of glutamatergic and GABAergic neuron type-specific exons with inferred positive and negative regulatory RBPs indicated on the right as red or blue tick marks, respectively. **F**, Similar to (E) but for CGE and MGE-specific exons.

To validate the inferred network, we first compared the predicted regulons to integrative splicing-regulatory network models of several well-studied neuronal RBP families, which were derived from analysis of multimodal datasets, including protein-RNA interactions and RBP-dependent splicing changes^41–43^. The target exon lists of ARACNe and the integrative models overlapped significantly (Figure 2A top panels). Importantly, among common exon targets identified by both methods, ARACNe predicted MOR typically agreed in the direction of regulation when compared to integrative modeling (Figure 2A bottom panels). This consistency was also seen between predicted regulons and exons showing altered splicing after RBP perturbation for several other RBPs without available integrative models^23,28,44^ (Figure 2B-C), including Elavl2, Elavl3, and Elavl4 (Figure 2B). The concordance of the different Elavl family members is notable since they display distinct expression patterns among neuron types (Figure S2), suggesting the capability of ARACNe to delineate specific targets regulated by unique members of a homologous RBP family.

**Figure 2.**
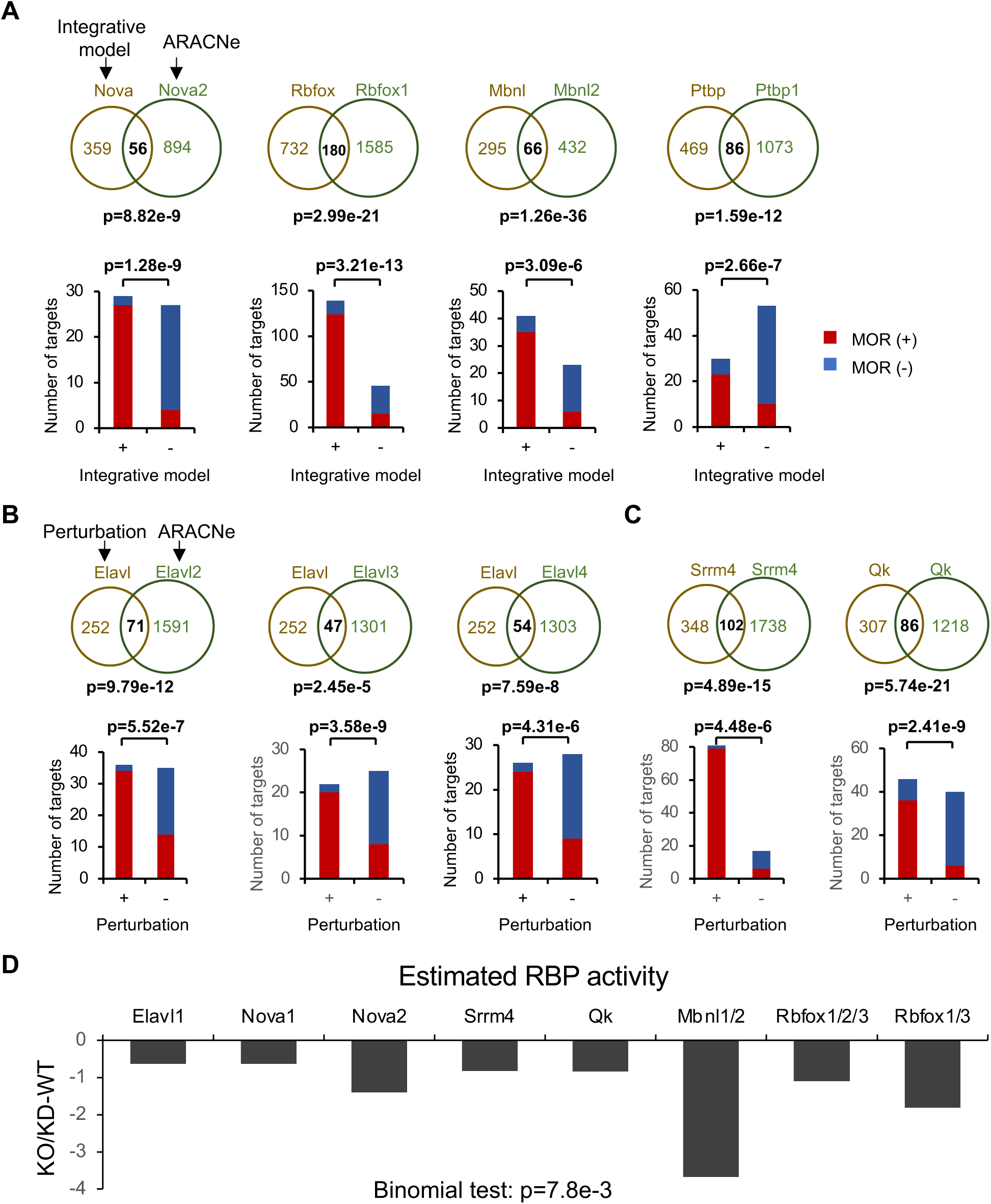
Validation of RBP regulons and activities inferred by ARACNe and VIPER algorithms. **A**, ARACNe-inferred regulons show significant overlap (Venn diagrams) with high-confidence lists of RBP family targets from integrative Bayesian models based on analysis of multimodal splicing data^44–46^. Among the exons common to both lists, the predicted directions, or modes of regulation (MOR), by VIPER are also highly consistent with the integrative models (bar graphs). P-values shown are derived from Fisher’s exact tests. **B,C**, Similar to (A) but additional regulons inferred by ARACNe were compared to RBP target lists identified from exon inclusion changes after RBP perturbations, as measured by RNA-seq or exon-junction microarray data. **D**, Bar plot of RBP activity difference estimates in RBP-depleted samples using the predicted regulons as compared to wild-type (WT) control. For perturbations of multiple RBP family members, activity value sums of the multiple RBPs perturbed were used. The statistical significance of the directional change was tested using a Binomial test.

RBPs have been shown to modulate tissue- or cell type-splicing in a binding position-dependent manner (aka, RNA map), with those binding upstream of target exons tending to repress the exon and those binding downstream tending to activate it^45–48^. Therefore, we visualized the distribution of RBP binding sites in the upstream and downstream introns of the target exons, as determined by bioinformatics motif predictions or RBP footprints mapped by crosslinking and immunoprecipitation (CLIP) tags. Consistent with direct position-dependent AS regulation, we found that exons predicted to have RBP-dependent inclusion (positive targets) tend to have motif and CLIP tag peaks downstream of the alternative exon and those predicted to have RBP-dependent exclusion (negative targets) tend to have peaks upstream of or inside the exon (Figure S3A-B, left panels). Notably, these patterns are also evident when only considering novel targets that were not previously identified by integrative network modeling (Figure S3A-B, right panels), suggesting that ARACNe predictions have identified previously unknown AS regulatory relationships.

We then estimated cell type-specific RBP activity from the inferred regulons using the VIPER algorithm (Table S2). Estimated RBP activity across neuron types is correlated with RBP expression level, as one would expect. However, the extent of correlation varies, likely because mutual information used in ARACNe captures both linear and nonlinear regulatory relationships (Figure S4). To evaluate the reliability of the results, RBP activity was estimated using various independent RNA-seq datasets comparing RBP knockout or knockdown samples and their controls^19,20,43,49,50^. This analysis confirmed that the inferred RBP activity was consistently lower in RBP depletion versus control samples (Figure 2D). In most cases we examined, the perturbed RBP was among those with the largest estimated activity reduction between the two compared conditions (Figure S5). These results suggest the activity metric captures *bona fide* differences in RBP activity levels.

### RBP activity estimations reveal candidate drivers of neuron type-specific AS programs

RBP activity estimations on the cell-type level allow the exploration of candidate RBPs mediating AS differences between various groups of cell types by performing differential activity tests (Table S3). As expected, known neuron-specific splicing factor RBPs tended to have higher activity in neurons versus glia (Rbfox1/2/3, Elavl2/3/4, Mbnl2, Khdrbs2/3 and Celf1/2/6), and the opposite was true for RBPs enriched in non-neuronal cells (Ptbp1, Srsf2, Srsf5, Hnrnpl, Qk; Figure 3A). Importantly, differential activity tests between clades of neuron types identified known and novel putative neuron class- and subclass-specific splicing factors. RBPs with the largest activity differences between the glutamatergic and GABAergic neuron classes included those we had previously identified based on their expression levels^15^ (Mbnl2, Celf1/2, and Khdrbs3 for glutamatergic, Elavl2 and Qk for GABAergic) as well as some previously unknown putative type-specific RBPs. These include Rbfox2/3 and Khdrbs2 for glutamatergic neurons and Elavl3 and Celf6 for GABAergic neurons (Figure 3B). Comparing the two GABAergic interneuron subclasses, we identified Elavl2 and Rbfox1 as candidates with MGE lineage-specific activity and Khdrbs2 and Celf1 as candidates with CGE-specific activity (Figure 3C).

**Figure 3.**
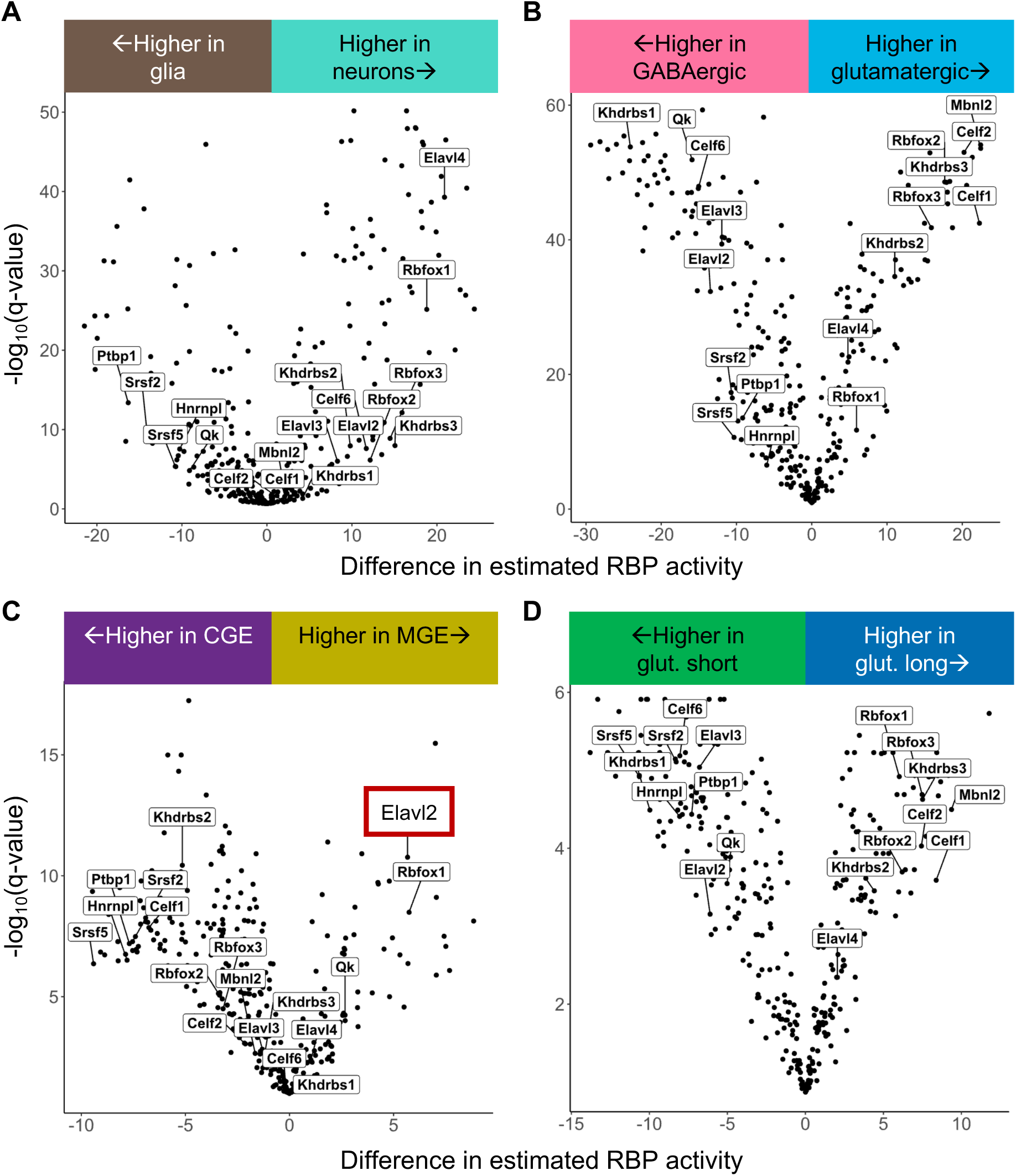
Differential RBP activity analysis provides candidate regulators of neuron type-specific splicing across different clades. In each panel, RBP activity was compared between two groups of cell types. **A**, neurons and glia. **B**, glutamatergic and GABAergic neurons. **C**, MGE- and CGE-lineage interneurons. **D**, long- and short-projecting glutamatergic neurons. Q-values are derived from empirical Bayes-moderated t-tests followed by multiple test correction.

We previously identified a group of exons showing differential splicing between neurons with long versus short-range projections^15^. To identify potential regulators of this AS program, we performed differential RBP activity tests between long-versus short-projecting glutamatergic neurons. This analysis revealed distinct sets of RBPs with higher activity in long-projecting glutamatergic neurons (Mbnl2, Celf1/2, Khdrbs2/3, and Rbfox1/2/3) and short-projecting glutamatergic neurons (Elavl2/3, Qk, and Celf6) (Figure 3D). Interestingly, these RBPs also show differential activities between glutamatergic and GABAergic neurons, which also differ in their range of projection.

Elavl2 modulates splicing predicted targeted exons in MGE-but not CGE-lineage interneurons. Although transcription factors controlling MGE-versus CGE-specific gene expression have been identified^9,10,51^, RBPs regulating the AS differences between these lineages are still unknown. To demonstrate the utility of MR-AS to generate testable hypotheses on AS regulatory programs in a specific cellular context, we decided to focus on Elavl2 (also known as HuB), which was predicted to have higher activity in MGE-than CGE-lineage interneurons (Figure 3C), consistent with its expression pattern from an early developmental stage^8,10,24^ (Figure 4A).

**Figure 4.**
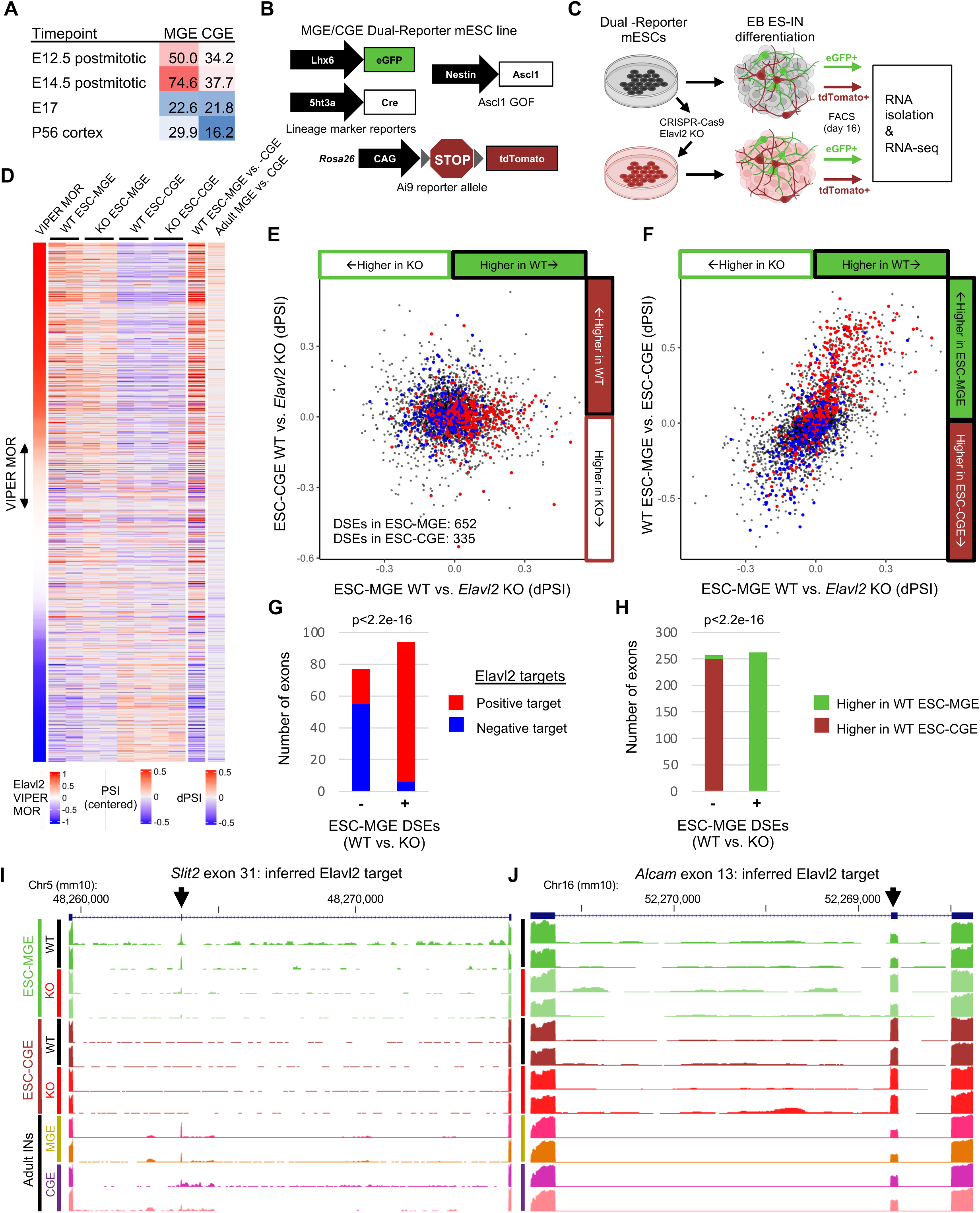
Validation of Elavl2 as a key MGE-lineage-specific splicing factor. **A**, *Elavl2* expression RPKM (reads per kb per million) in MGE- and CGE-lineage interneurons at various time points during cortical development. The gene shows a consistent preferential expression in MGE-lineage neurons as early as embryonic day 12.5 (E12.5) and persists into adulthood. **B**, Construct design for the MGE/CGE dual-reporter mouse ESC line. eGFP is positioned downstream of the MGE-specific *Lhx6* promoter, and tdTomato is contained in an Ai9 reporter with Cre under the control of the CGE-specific marker *5ht3a*. Ascl1/Mash1 overexpression is driven by the neural progenitor marker *Nestin* to promote interneuron differentiation. **C**, Experimental schema for testing the role of Elavl2 in interneuron type-specific splicing regulation. *Elavl2* was knocked out in the dual-reporter mouse ESC line using CRISPR-Cas9. WT and KO ESCs were differentiated into interneurons using an embryoid body-based protocol. On day 16 of differentiation, cells were isolated based on reporter fluorescence by FACS and RNA was isolated for RNA-seq. **D**, Centered exon inclusion (percent spliced in or PSI) values of inferred *Elavl2* targets ordered by predicted MOR. Inclusion differences of these exons in WT GFP+ (ESC-MGE cells) vs. tdTomato+ (ESC-CGE cells) samples and adult MGE-vs. CGE-lineage interneurons are shown at right. **E-F** Scatter plots (top) and quantification of directionality agreement (bottom) for predicted *Elavl2* target inclusion value differences in WT versus *Elavl2* KO eGFP+ samples (x-axis) compared to WT versus *Elavl2* KO tdTomato+ samples (y-axis, e) or WT eGFP versus WT tdTomato+ samples (y-axis, F). Positive and negative *Elavl2* targets predicted by ARACNe/VIPER are indicated in the scatter plots by red and blue, respectively. P-values above the barplots are calculated by Fisher’s exact test. **G,H**, Genome browser views of *Slit2* exon 31 (G) and *Alcam* exon 13 (H) inclusion in WT or KO ESC-interneurons and adult MGE- or CGE-lineage neurons. Both exons are inferred targets of *Elavl2*.

To confirm the predicted MGE-specific Elavl2 activity, we utilized a mouse embryonic stem cell (ESC) line with two fluorescent reporter constructs to mark the MGE- and CGE-lineage cells when they are differentiated into interneurons (ESC-MGE and ESC-CGE) (Figure 4B-C). The construct contained *eGFP* downstream of the *Lhx6* promoter, which is specifically expressed in MGE-lineage cells, and *tdTomato* under the control of an Ai9 reporter allele driven by *5ht3a*, a CGE-lineage marker. To encourage neuronal differentiation, an *Ascl1* (encoding Mash1) gain-of-function construct driven by *Nestin* was also incorporated (Figure 4B). We used CRISPR-Cas9 genomic engineering to generate a homozygous *Elavl2* knockout (KO) line from this parent line (Figure 4C, Figure S6 and Methods). Wild-type (WT) and *Elavl2* KO lines were differentiated into ESC-interneurons using an established embryoid body-based differentiation protocol^52,53^. We then isolated the differentiated neurons using fluorescent activated cell sorting (FACS), collecting eGFP+ ESC-MGE and tdTomato+ ESC-CGE populations from both genotypes for RNA isolation, RT-qPCR and RNA-seq (Figure 4C and Figure S7A-B). ESC-MGE and ESC-CGE WT and KO samples were enriched in previously known MGE- and CGE-lineage marker expression^8,51^, respectively (Figure S7C-E), indicating that the differentiated ESC-interneurons had committed to the appropriate developmental lineages. ESC-MGE cells also showed higher expression of some MGE-specific RBPs we identified in adult, including *Elavl2* (Figure S7F). Furthermore, although ESC-INs are immature, we observed differential splicing between ESC-MGE and ESC-CGE cells, in consistent directions as we observed in adult neocortex (Figures 4D and S8; Table S4), indicating that the differential splicing-regulatory programs between MGE- and CGE-interneurons *in vivo* is recapitulated in the *in vitro* experimental system.

When we examined the impact of Elavl2 on neuron type-specific splicing, we found that exons predicted to be activated by Elavl2 (with a positive MOR) show higher inclusion in ESC-MGE cells, while exons predicted to be repressed by Elavl2 (with a negative MOR) show higher inclusion in ESC-CGE cells (Figure 4D). Importantly, consistent with a predicted higher activity, *Elavl2* knockout caused splicing changes in a much larger number of exons in MGE-like cells than in CGE-like cells (707 vs. 379, FDR<0.05, |ΔΨ|≥0.1; Figure 4E top panel). Upon Elavl2 depletion in ESC-MGE samples, exons predicted to be activated by Elavl2 mostly showed more skipping (88/110=80%), while exons predicted to be repressed by Elavl2 showed more inclusion (55/61=90.2%; p<2.2e-16; Fisher’s exact test) (Figure 4G). Moreover, exons showing reduced exon inclusion upon Elavl2 depletion in general have a higher inclusion in ESC-MGE cells, while exons showing increased exon inclusion on Elavl2 depletion in general have a higher inclusion in ESC-CGE cells, which largely correspond to lineage-specific exons observable in adult neurons (Figure 4F and Figure S8). These data suggest that Elavl2 has a higher activity in the ESC-MGE cells than ESC-CGE cells and its depletion resulted in diminished differences between the two cell populations, supporting the hypothesis that Elavl2 plays a pivotal role in driving AS differences in MGE-versus CGE-lineage interneurons.

MGE- and CGE-lineage neurons differ in the migratory path they take during development and settle into different distributions among the cortical layers^54^. Exons predicted to be Elavl2 targets include those in genes that have roles in neuronal migration (Table S5). Among these, *Slit2* exon 31, a predicted target of Elavl2, had much lower inclusion in KO than WT ESC-MGE samples, in agreement with higher inclusion in adult MGE-lineage interneurons regulated by Elavl2 (Figure 4G). *Slit2* is part of the Robo-Slit pathway, which is critical for axon guidance and cell migration during development^55–57^. Another example, exon 13 of the *Alcam* gene, which encodes a cell adhesion molecule involved in migration, neurite extension, and axon guidance^57,58^, showed the opposite pattern of inclusion, consistent with its predicted negative regulation by Elavl2 in adult MGE-lineage interneurons (Figure 4H). In dictating MGE-specific splicing programs early in development, Elavl2 may have a role in establishing distinct migratory properties of MGE- and CGE-lineage interneurons in accordance with the needs of each lineage.

### Identification and validation of a splicing program differentially regulated in long-versus short-projecting neurons

As a second case study of MR-AS applications, we decided to test the impact of neuron type-specific AS regulation on neuronal morphology, specifically the range of axonal projections. In a recent study, we identified a subset of exons showing differential splicing between long-vs. short-range projecting neurons across multiple clades^15^. However, their functional significance has not been validated and the upstream regulators that drive differential AS are unknown. To address these questions, we first refined our analysis to identify a common module of 61 cassette exons that show differential inclusion between long- and short-range projecting neurons.

The exons are differentially spliced between glutamatergic and GABAergic neurons (which mostly have long and short projections, respectively), long- and short-projecting glutamatergic neurons, and long- and short-projecting Somatostatin (Sst)-positive interneuron types (Figure 5A-B and Table S6). The genes harboring these exons are enriched in gene ontology (GO) annotations related to the establishment or maintenance of axons including “cell projection organization” and cellular component terms related to axonal projections, including “presynapse”, “cell projection”, and “distal axon” (Figure 5C and Table S7).

**Figure 5.**
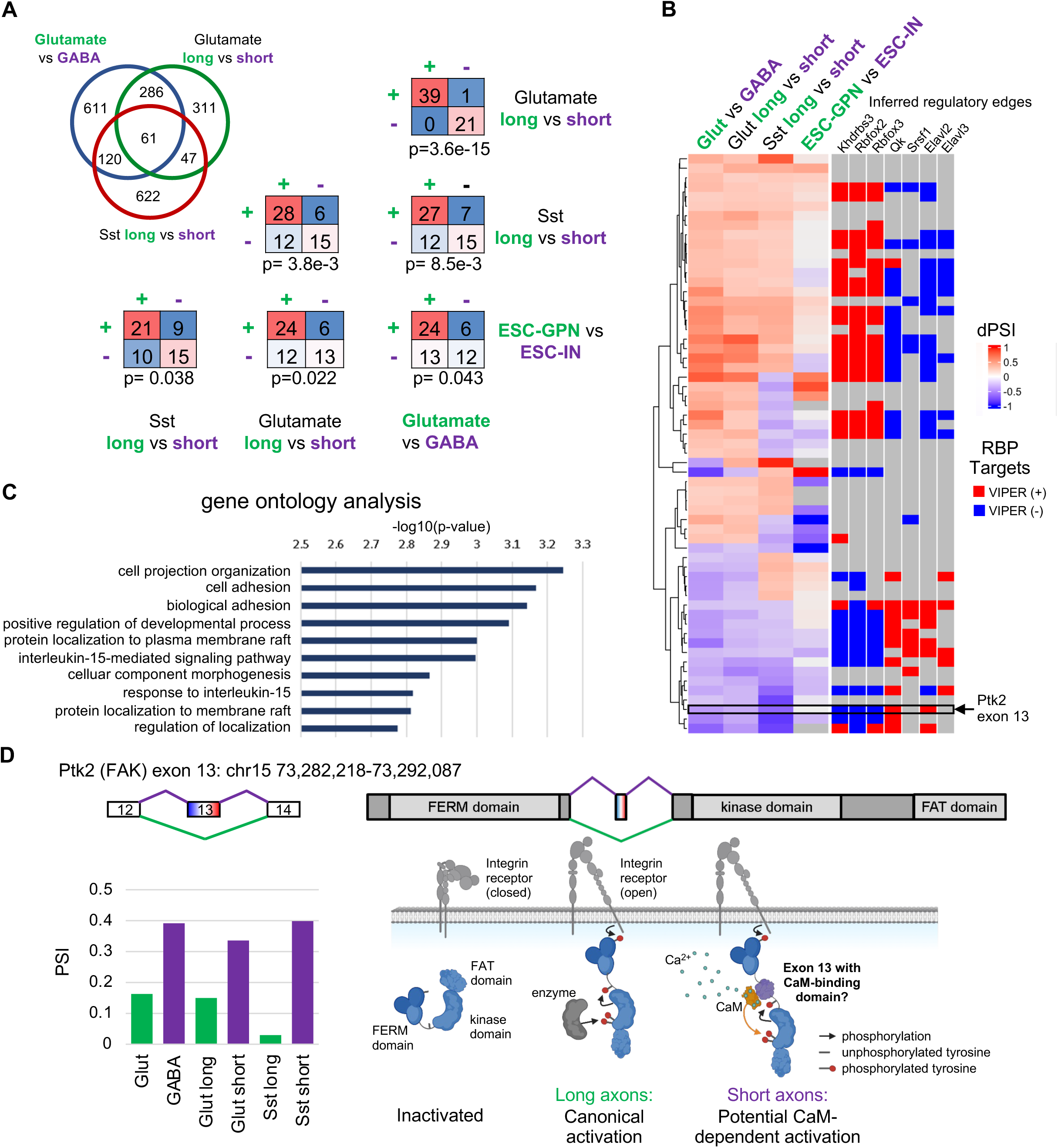
Characterization and validation of an exon module specific to long- or short-projecting neurons. **A**, Top left: Venn diagram showing the overlap of exons differentially spliced between glutamatergic versus GABAergic neurons, long-versus short-projecting glutamatergic neurons, and long-versus short-projecting Sst interneurons. Bottom right: Comparisons in the directionality of differentially spliced exons in the identified common exon between each of the neuron type comparisons, including a comparison of ES-derived globus pallidus neurons (ES-GPNs) and ES-derived interneurons (ESC-INs). P-values are calculated by Fisher’s exact test. **B**, A heatmap showing the differential inclusion of the 61 exons identified in all three comparisons. Inferred positive (red) ore negative (blue) regulators are indicated on the right. **C**, Gene ontology (GO) analysis of genes containing the 61 overlap exons shows an enrichment of genes related to axonogenesis and their maintenance. **D**, A schematic depicting *Ptk2* exon 13, which is specifically included in short axon neurons. The cartoon depicts the possible biological function of the exon, which contains a reverse calmodulin binding domain and may link the gene’s function to calcium signaling.

The majority of the exons show consistent preferences between the comparisons for exon inclusion in long- or short-projection neurons (p=3.6e-15, p=3.8e-3, and p=8.5e-3 by Fisher’s exact tests; Figure 5A, top three tables). Some reside in genes that suggest a long- or short-projecting neuron type-specific tailoring of calcium-responsive gene function, such as those in calmodulin-dependent serine kinase II delta (*Camk2d*) and calmodulin-dependent serine protein kinase (*Cask*, data now shown). A particularly interesting example is exon 13 of *Ptk2* (also known as focal adhesion kinase or FAK), which shows a strong preference for inclusion in short-projecting compared to long-projecting neuron types (Figure 5D). This alternative exon contains a reverse calmodulin (CaM) consensus sequence, a motif involved in binding CaM similarly to the forward sequence^59,60^. The differential Inclusion of this exon raises the possibility that FAK signaling is modulated to allow activation by CaM in short-projecting neurons (Figure 5D). Among its many known roles in axon outgrowth, FAK has already been shown to modulate axonal branching in a calcium/CaM-dependent manner in hippocampal neurons^61^. It is possible that this mode of signaling is carried out by FAK isoforms containing exon 13, which are enriched in short-projecting neuron types.

To validate the functional impact of the AS module on neuronal projection, we utilized comparison of ESC-interneurons and ESC-derived globus pallidus neurons (ESC-GPN), modeling an intriguing pair of cell lineages that share a common developmental origin in the MGE but diverge into short-projecting cortical interneurons and long-projecting globus pallidus midbrain neurons, respectively. We recently demonstrated that while overexpressing the transcription factors *Nkx2.1* and *Dlx2* in ES cells promotes interneuron differentiation, adding the overexpression of *St18* is sufficient to drive them to acquire a globus pallidus-like state, including long and elaborated axons^62^. We leveraged this system to investigate whether differential AS regulation may covary with neuronal projection length by examining the state of the putative long- or short-projection-specific AS module using RNA-seq data from the two cell populations. Importantly, we found that these exons showed consistent splicing switches in reprogrammed ESC-GPNs as compared to ESC-interneurons, as one would expect from the splicing differences between the other types of long- and short-range projecting neurons. Exons with increased inclusion in ESC-GPN are most frequently more included in glutamatergic neurons as compared to GABAergic neurons, as well as in long-range projecting glutamatergic and GABAergic neurons than their counterparts with short-range projection (p<0.05 in all comparisons by Fisher’s exact tests; Figure 5A-B). This result provides an independent line of evidence that the identified splicing module is likely important for tailoring transcriptomes to the needs of establishing or maintaining long- or short-projecting neurons across diverse neuronal classes.

We next examined possible RBP splicing regulators of the long- or short-projection-specific AS module by comparing the estimated RBP activity between each of the long-versus short-projection neuron types. Intriguingly, principal component analysis (PCA) of estimated neuron-type specific RBP activity placed near-projecting glutamatergic and long-projecting Sst neurons in an intermediate state between those of the majority of the glutamatergic and the majority of the GABAergic neuron types along PC1, indicating large differences in RBP activity along this axis (Figures 6A-B and S9). RBP activity differences between each of the long-versus short-projecting neuron comparisons revealed a number of RBPs with consistent preference for a projection type, including Rbfox3, Khdrbs2, and Khdrbs3 for long-projecting neuron-specific activity and Qk, Srsf1, Elavl2, and Elavl3 for short-projecting neuron-specific activity (Figures 6C-E and S9). Differential inclusion levels of exons in the long- or short-projection-specific module contained in the regulons of these RBPs confirm that the directional differences between the long-versus short-projection neuron types are consistent with the exons’ predicted direction of regulation by the RBPs. For example, exons activated by Rbfox3 and Khdrbs3 in general showed higher inclusion in long-projecting neurons, while exons activated by Qk and Elavl2 showed higher inclusion in short-projecting neurons; the opposite patterns were seen for exons repressed by these RBPs (Figure 6F-G). These data suggest that neuron type-specific activity of these RBPs play a role in driving the differential splicing programs. This example again demonstrates the power of the inferred splicing-regulatory networks and RBP activity estimations to discover functionally relevant exon modules and the underlying regulatory mechanisms.

**Figure 6.**
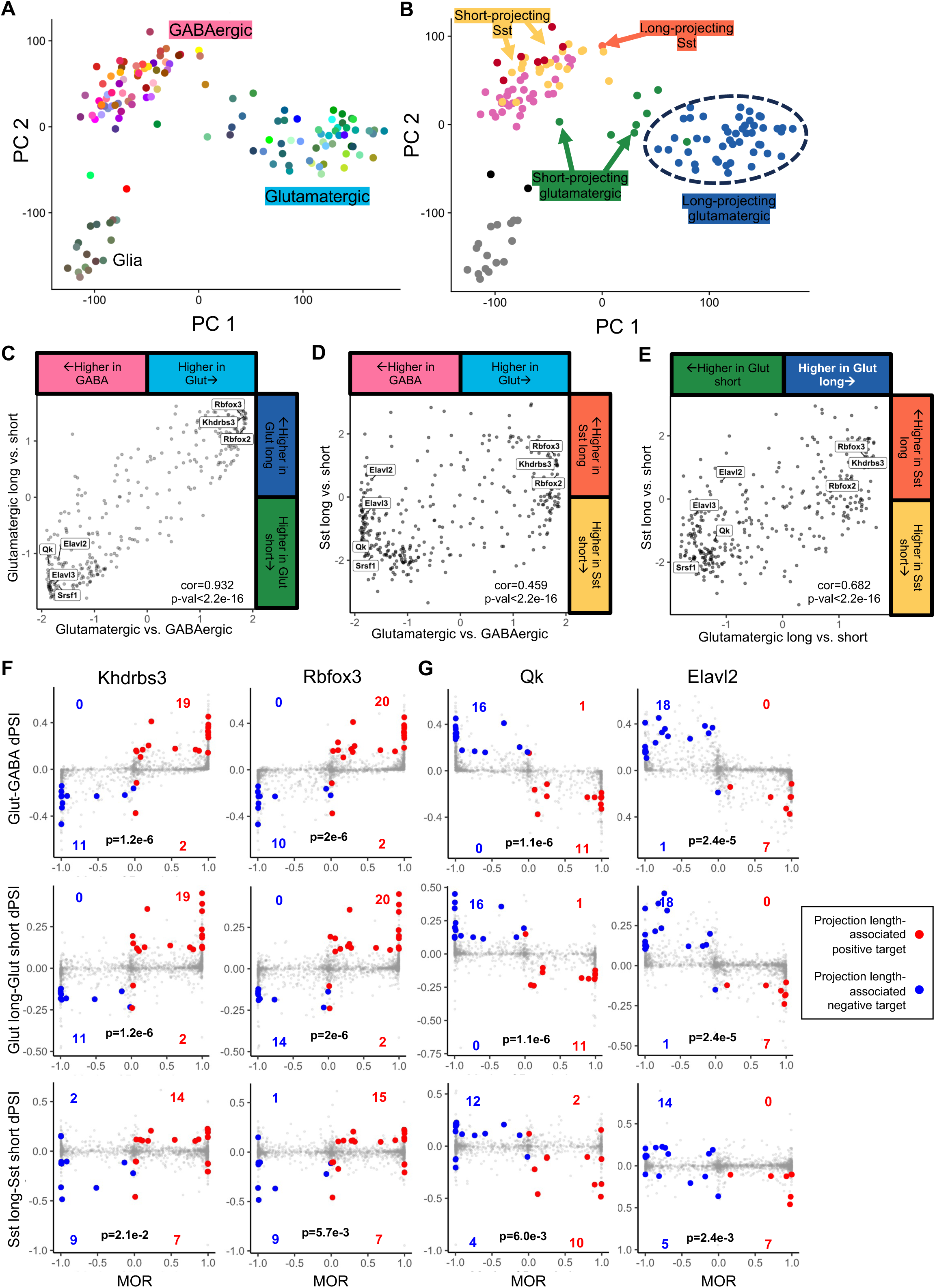
Inferred networks identify candidate drivers of projection length-associated splicing. **A**, PCA plot of the first two principal components of RBP activity with cell type clusters and neuronal classes indicated by color. **B**, Similar to (A) but with neuron projection types indicated by color. **C-E**, Comparisons of differential RBP activity in long-versus short-axon neuron types correlate across different neuronal classes (Pearson). **F-G**, Conchordance of candidate RBP regulons with the module of projection-length associated exons identified in Figure 5. Each scatter plot shows the predicted mode of regulation by the RBP on the x-axis and change in exon inclusion between projection types on the y-axis. Positive and negative target exons overlapping with the projection length-associated module and are differentially spliced between the groups are colored and counted in red or blue, respectively, and all other exons in the regulon are shown with reduced opacity. P-values are from Fisher’s exact test.

## Discussion

Recent efforts on transcriptome profiling using deep sequencing have revealed an enormous diversity of AS among brain cell types^13,15,24,32,34,63,64^, but technical and analytical challenges have limited the pace of progress elucidating the regulatory programs driving these patterns. Previously, splicing-regulatory networks are typically inferred by focusing on the regulons of individual RBPs, which can be interrogated using multifaced assays, such as mapping RBP-binding sites by CLIP or bioinformatic predictions of motif sites, and identifying exons with altered splicing upon perturbation of RBPs. These datasets can then be integrated, e.g., using Bayesian networks, to define RBP regulons with confidence^41–43^. Together with cell type-specific RBP expression and exon splicing patterns, these regulons allowed us to evaluate glutamatergic or GABAergic neuron-specific activity for a select of well-studied RBPs^15^. Despite the high accuracy of this approach, it involves substantial efforts to produce the required datasets for network inference, which is challenging to scale. Furthermore, experiments are also frequently performed in bulk tissues so that cell type specificity of the regulons is difficult to delineate.

This study presents an alternative strategy to infer splicing-regulatory networks by correlating exon inclusion and RBP expression across a large panel (>100) of cell types with rich regulatory dynamics and then estimate neuron type-specific RBP activity levels based on aggregated splicing patterns of the regulon for each RBP. The advantage of this strategy is that it requires only expression-profiling data, which are nowadays routinely obtained with advancements in deep sequencing technologies. The effectiveness of this strategy for reverse engineering of transcriptional regulatory networks and master regulator analysis has been well established^37,39,65,66^. With the availability of similar scRNA-seq datasets that can simultaneously quantify RBP expression and exon splicing across over 100 neuronal types, here we demonstrate that this strategy, as implemented in MR-AS, can also be used to systematically reverse engineer splicing-regulatory networks and identify driving factors of neuron type-specific splicing regulatory programs. We present a comprehensive network composed of 8,336 cassette exons and 350 RBPs connected by 174,274 edges inferred from neocortical cell types in the adult mouse brain, with an average of 498 target exons per RBP, together with neuron type-specific activities for each RBP.

Multiple lines of evidence support the validity of the inferred cell splicing-regulatory network and RBP activities. First, the inferred RBP targets overlap significantly with high-confidence target lists defined by integrative modeling using independent, multimodal datasets. Importantly the overlapping exons agree very well in the inferred direction of regulation (Figure 2A-C). Second, inferred RBP targets have enriched binding sites as evidenced in CLIP and motif data in positions consistent with the direction of splicing regulation. In addition, the inferred regulons by ARACNe include *bona fide* targets not previously identified by the integrative modeling approach (Figure S3). These novel targets could have been missed previously due to various reasons such as dilution of cell-type specific regulatory effects in bulk tissues used in previous analyses. Third, as one would expect, the estimated RBP activity values are consistently lower in the RBP knockout samples than their wild-type counterparts using independent datasets (Fig 2D). These data suggest that our analyses provide a reliable resource to generate testable hypotheses of neuron type-specific AS.

This method allowed us to identify novel candidate regulators of neuron type-specific splicing for well-defined neuronal subclasses based on their estimated RBP activity differences. To validate our predictions, we focused on Elavl2 showing MGE-lineage interneuron-specific activity. The Elav-like genes encode a highly conserved RBP family that have emerged as critical regulators of neural development and function. Three of the four family members, Elavl2, Elavl3, and Elavl4 (also known as HuB, HuC, and HuD), are selectively expressed in neurons early and throughout neuronal development, and they function as important splicing factors for proper neuronal differentiation and synaptic plasticity^67–72^. However, specific functions of individual factors on neuronal identity have not been elucidated. *Elavl2*, a risk gene for schizophrenia^73^, is expressed on a different timeline and in different cell types compared to *Elavl3* and *Elavl4*. During neurogenesis, Elavl2 is expressed in neurons immediately after their birth in the ventricular zone, whereas Elavl3 and Elavl4 are expressed at later timepoints^67,74^. In the hippocampus, *Elavl3* and *Elavl4* show broad expression amongst neuron types while *Elavl2* is specifically expressed in CA3 pyramidal neurons and hilar interneurons^75^. These observations imply potential functional differences among Elavl family members in different neuron types, supporting our hypothesis on the role of Elavl2 in MGE-lineage interneurons. Indeed, when we abrogated the expression of *Elavl2* in an ES-derived model of MGE- and CGE-lineage interneurons, predicted Elavl2 targets showed specific splicing changes in ESC-MGE neurons, as compared to ESC-CGE neurons so that differential splicing between the two cell populations became diminished, confirming our hypothesis. These results validate the utility of MR-AS in identifying candidate RBPs driving cell type-specific AS delineating canonical neuronal types.

Epigenetic and transcriptional regulation have been repeatedly demonstrated to play a dominant role in specifying developmental cell lineages^9,51,54,76–80^. Intriguingly, while much of splicing varies by tissue in concordance with gene expression at the transcriptional level^81–83^, it has been a longstanding observation that tissue- or cell type-specific genes are frequently regulated at the transcriptional and splicing levels in distinct patterns^19,83,84^, raising the possibility that AS provides an orthogonal regulatory mechanism to specify specific neuronal properties shared across developmental lineages^15^. AS is uniquely situated to modulate genetic programs in response to various signaling inputs since post-transcriptional processing occurs at a shorter timescale compared to transcriptional regulation. For example, multiple neuronal RBPs have been shown to modulate AS in response to synaptic activity^85–88^. The neuronal projection-specific AS module we identified across several disparate neuronal clades is another possible example of orthogonal AS regulation to meet biological needs of particular cell types or states shared across different developmental lineages. We note that the identification of the long- or short-projection neuron-specific AS program required a wide diversity of cortical neurons to be sampled in both glutamatergic and GABAergic lineages for systematic comparison, and its generality would have been missed in an analysis using a more limited number of cell types. As groups of long- and short-projection neuron types are found separately within the larger classes of glutamatergic neurons and Sst-positive GABAergic interneurons, traditional lineage marker-based isolation of the groups to examine their AS differences would be a tedious and resource-intensive process. The MR-AS framework offers unique flexibility to compare groups of neuron types that differ in certain functional properties, regardless of their developmental lineages, and generate testable hypothesis of the underlying molecular mechanisms.

Altogether, the differential AS programs we inferred between canonical neuronal subclasses and within multiple unrelated classes illustrate the power of an unbiased, comprehensive mapping of cell type-specific splicing-regulatory networks. As ever larger and higher-resolution scRNA-seq data sets become available, we expect such analyses will likely reveal AS regulatory programs both parallel and orthogonal to the transcriptional cell type hierarchy, a key aspect to understanding neuronal function in physiological and pathological conditions.

## Methods

### RNA-seq data preprocessing

Quantification of cassette exon inclusion and RBP expression using adult cortex scRNA-seq data was obtained from our previous study^15^. Briefly, scRNA-seq reads^8^ were mapped to the mm10 genome using OLego^89^. Gene expression and exon inclusion values were quantified using the Quantas pipeline (https://zhanglab.c2b2.columbia.edu/index.php/Quantas) as previously described^43,91^. In this study, we quantified exon inclusion and gene expression for 133 cell clusters (cell types) as defined by the original study authors^8^.

Aligned reads from single cells were pooled per cluster before quantification. To be included in network inference using ARACNe, we require exons to be quantified in ≥ 25 clusters, and for the remaining 12903 exons after this filtering, missing values of exon inclusion were imputed by knn (k=10). For RBPs, we used 376 genes from RBPdb (http://rbpdb.ccbr.utoronto.ca/), supplemented by literature search that identified ∼20 additional RBPs not included in RBPdb, such as *Srrm4*/nSR100. RNA-seq data of ESC-interneurons and ESC-GPNs were obtained from a previous study^62^ (SRA accession: SRP329886) and analyzed similarly using the Quantas pipeline.

### ARACNe and regulon inference

Gene expression data have been used to reconstruct transcriptional regulatory networks in various biological contexts. ARACNe (algorithm for the reconstruction of accurate cellular networks)^37,38^ and its companion algorithm VIPER (virtual inference of protein activity by enriched regulon analysis)^39,40^ uses mutual information estimated across diverse cellular conditions to associate regulators with their targets, whose collective behavior in a sample is then used to estimate the regulator’s activity. This approach has the advantage of aggregating large numbers of targets to estimate a regulator protein’s activity rather than relying on single gene expression values of the regulators, which can be an unreliable approximation of activity. While it has been successfully used to infer the downstream target networks of transcription factors and signaling molecules and estimate cell state-specific regulator activity, this method has not been used to model AS regulation, which has to be optimized due to different noise structures in the scRNA-seq data at the splicing level.

We used ARACNe-AP^37,65^ to infer splicing-regulatory networks at the neuron type (cluster) level based on the mutual information of RBP expression and target exon inclusion with several modifications. First, we did not apply data processing inequality (DPI) to eliminate potential indirect regulations. This is because estimation of mutual information is noisier for scRNA-seq data compared to bulk RNAseq. Importantly, estimation of gene expression and exon splicing is subject to different levels of noise, which complicates DPI assessment. Second, we examined a set of well-studied RBPs to calibrate the number of targets predicted by ARACNe using varying mutual information thresholds. The number tends to vary dramatically for different RBPs. For results presented in the paper, we set mutual information p-value = 1e-8, while limiting the maximum number of targets per RBP to be 1000 for each bootstrapped network. We then consolidated 100 bootstrapped networks to keep only reproducible edges by Bonferroni-corrected Poisson p-value=0.05. This resulted in a total of 174,274 edges in the inferred regulatory network between 350 RBPs and 8,336 cassette exons.

After the regulatory network was derived, we used the aracne2regulon implemented in the VIPER package to infer the mode of regulation (MOR, i.e., the probability of a target being activated or repressed by an RBP). MOR is calculated based on the Spearman rank correlation between the expression of an RBP and the inclusion of its target exons. A 3-component Gaussian mixture (representing activation, with positive correlation, repression, with negative correlation, and uncertain direction, with correlation around zero) was then used to infer MOR. The mutual information was recorded to represent the likelihood of regulation.

In addition to cluster-level analysis, we also inferred the regulatory network by correlating RBP expression and exon inclusion at the single-cell level. We noticed that the regulatory network and MOR inferred at the cluster level provided the most accurate results, as judged by comparison with targets derived by integrative modeling or perturbation experiments^41–43^. Therefore, for the results presented in this study, we used the network inferred at the cell cluster level.

### Inference of RBP activity

We estimated RBP protein activity across 133 neuron types using regulons (i.e., regulatory networks and MOR) inferred at the neuron type level. For this analysis, the exon inclusion level was standardized to zero mean and unit variance across cell types. VIPER analysis was performed without pleiotropy correction, which was designed to punish enrichment of targets that are co-regulated by other regulators, since this feature led to an overly conservative set of inferred RBP targets.

### Gene ontology analysis

Genes containing exons of interest were tested for gene ontology (GO) term enrichment using the R package “topGO”. GO terms with a Bonferroni-corrected Fisher’s test p-value of ≤0.05 were shown in the figures.

### MGE an CGE dual reporter ESC line construction

To derive the MGE and CGE dual-reporter ESC line, super ovulated 5HT3aR-BACCRE/+; Ai14/Ai14 (MMRRC:036680-UCD; JAX 007909) females were mated with Lhx6-eGFP (MMRRC 000246-MU) males, and late-stage blastocysts were harvested and cultured for outgrowth following established protocols^92^. Genomic DNA isolated from generated lines were genotyped for Cre, Ai14, or eGFP according to genotyping protocols for corresponding mouse lines. Lines with all alleles were further propagated and tested for faithful recapitulation of reporter expression following differentiation. Nestin-Ascl1-ires-tTA22; TRE-Dlx2 (NAIT) constructs were cloned by modification of constructs previously published^52^ using standard methods and introduced into the dual reporter ESC background by co-nucleofection with puromycin selection cassette and antibiotic selection, clonal isolation, and genotyping.

### CRISPR/Cas9 genome engineering

*Elavl2* knockout synthetic guide RNAs (sgRNA; Synthego multi-guide CRISPR Gene Knockout Kit v2) and Alt-R S.p. HiFi Cas9 Nuclease V3 protein (IDT, 1081058) were introduced into low-passage dual-reporter ESCs using the Mouse Embyronic Stem Cell Nucleofector Kit (Lonza VPH-1001). ESCs were treated with 0.025% trypsin/1% EDTA (Gibco) to form a single cell suspension in the appropriate kit reagents, electroporated with sgRNAs and Cas9 protein (Lonza Nucleofector, program A-24), and replated into 24 well plates. After 6 days in culture, they were sorted into 96 well plates at clonal density. Individual clones were then expanded for genotyping and screening for *Elavl2* knockout by Sanger sequencing. Two clones containing the homozygous, frameshifting *Elavl2* knockout were identified and expanded over the course of 4-5 passages before downstream experiments.

### ESC culture and neuronal differentiation

Mouse ESCs were cultured for at least two passages on 0.1% gelatin and fed every 1-2 days with ESC medium consisting of DMEM (EMD Millipore SLM220B) supplemented with ESC-screened FBS (Hyclone SH30071.03E), 1x Modified Eagle Medium Non-Essential Amino Acids (Gibco), 1x Sodium Pyruvate (Gibco), 1x Glutamax (Gibco), 1x Penicillin/Streptomycin (Gibco) , 10 uM beta-mercaptoethanol (Fisher Scientific), and 10^4^ U/mL ESGRO LIF (Sigma Millipore). ESCs were differentiated into ESC-interneurons as described previously^54,69^. Briefly, ESCs were dissociated using 0.025% trypsin/1% EDTA (Gibco) and suspended at a density of 50,000 cells/well in non-TC-treated 24 well plates in 1mL of differentiation media consisting of Glasgow’s Modified Eagle Medium (Gibco), 1x Penicillin/Streptomycin (Gibco), 1x Modified Eagle Medium Non-Essential Amino Acids (Gibco), 1x Sodium Pyruvate (Gibco), and 1x Glutamax (Gibco). The Wnt inhibitor XAV-939 (Tocris, 1.5uM) was added to this initial suspension to promote a telencephalic neural fate. On day 4, medium was replaced by the same mixture with both XAV-939 and sonic hedgehog agonist (SAG; Tocris, 0.1uM) to ventralize the cells, taking care not to disturb the newly formed EBs. On day 6 and every 3 days until day 15, medium was replaced with the mixture with SAG but not XAV-939.

### ESC-interneuron FACS sorting, RNA collection and sequencing

On day 16 of ESC-interneuron differentiation, WT and *Elavl2* knockout EBs were collected into Eppendorf tubes where the medium was replaced with papain solution containing 20 U/mL papain and 67 U/mL DNase (Worthington). Tubes were incubated for 45-60 minutes at 37° C under constant agitation. Papain solution was replaced with diluted ovomucoid inhibitor mixture with 67 U/mL DNase and gently triturated 10 times using M937 syringe needle tips. The supernatant cell suspension was then removed from residual EB clumps and layered over non-diluted ovomucoid inhibitor mixture and centrifuged at 100g for 6 minutes. Cell pellets were then resuspended in FACS sort buffer consisting of DPBS with no calcium or magnesium supplemented with 20mM HEPES (Thermo Scientific), 5mM EDTA (Thermo Scientific), 20 U/mL DNase I (Worthington), and DAPI (Sigma-Aldrich). The suspension was passed through a 70 uM cell strainer, and the viability of the single cell suspensions were confirmed to be at least 95% viable using a Countess II cell counter (Thermo Scientific). These single cell suspensions were then FACS sorted into eGFP+ and tdTomato+ populations (Supplementary Figure 5) in differentiation medium supplemented with 5% FBS, using standard gating to remove debris, doublets, and nonviable cells. The collected cells were kept on ice and then resuspended in Trizol for RNA isolation. Samples were processed using either the standard Trizol (Thermo Scientific) protocol or using an RNeasy Micro Kit (Qiagen) using the aqueous RNA phase from samples centrifuged in Trizol. Isolated RNA was evaluated using a Bioanalyzer to confirm that RNA integrity numbers (RIN) were above 8. High-quality samples were submitted for RNA sequencing at the Columbia Genome Center, using TruSeq chemistry (Illumina) with poly-dT selection. Samples were multiplexed in each lane of a NovaSeq 6000 (Illumina) and 40 million 2×100 bp paired-end reads were generated per sample using RTA (Illumina) for base calling. RNA data were processed and analyzed as described above.

## Data and code availability

Illumina sequencing data from the Elavl2 WT and KO ESC-MGE and ESC-CGE cells will be deposited to NCBI Short Read Archive (SRA). The MR-AS software package is available at https://github.com/chaolinzhanglab/mras.

## Acknowledgements

We thank Zhang lab members and Vilas Menon for helpful discussion throughout the project. This study was supported by grants from the National Institutes of Health (NIH) (K99NS121275 to MC, R35A197745 to AC, R01NS117695 to EA, R01NS125018 and R35GM145279 to CZ). DFM was supported in part by NIH training grant T32NS064928. This research used the Genomics Shared Resource and CCTI Flow Cytometry Core, as supported in part by NIH awards S10RR027050 and S10OD020056, and P30CA01369.

