## Supplemental Figures S1-S9 for "Reverse engineering neuron type-specific and type-orthogonal splicing-regulatory networks using single-cell transcriptomes"

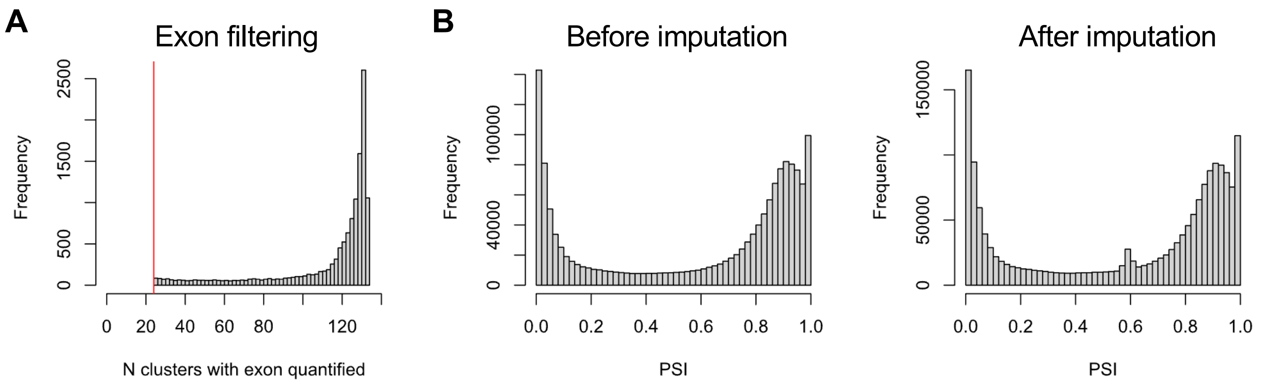


**Figure S1. Exon filtering and missing value imputation. A**, Exons were filtered by requiring splicing quantifiable in ≥25 cell types (clusters) with ≥ 20 junction reads. This filter resulted in 12,903 exons used for network inference. **B**, Distributions of PSI values before and after imputation of missing values using K-nearest neighbor (knn, k=10) estimation. Note the overall distribution of PSI values preserve after imputation despite the increase in exon numbers shown in y-axis.

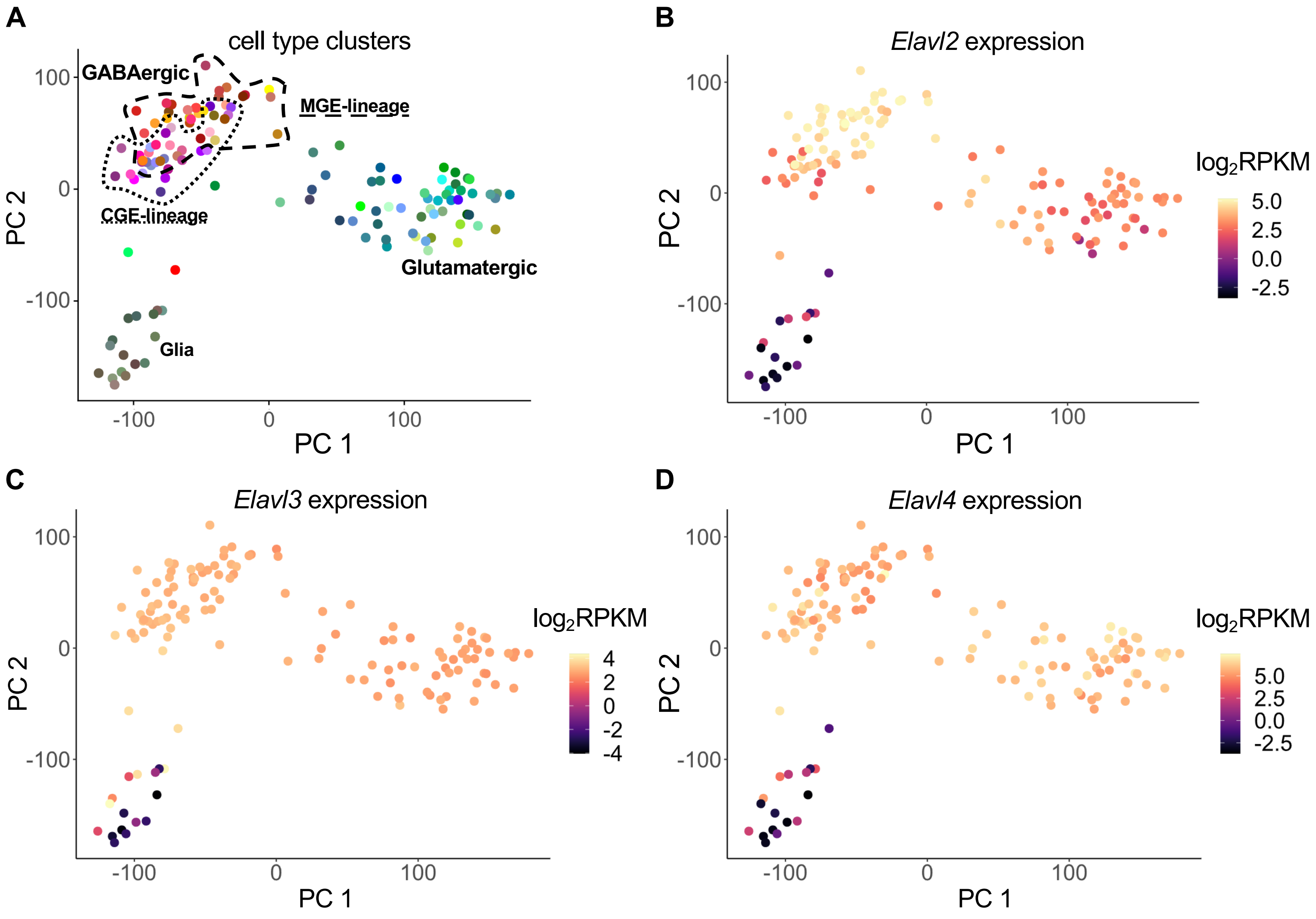


**Figure S2. Differential gene expression patterns among Elavl family RBPs. A**, PCA plot of RBP activity with cortical cell type clusters indicated by color. **B-D**, Similar to (A) but with cell type clusters colored by *Elavl2* (B), *Elavl3* (C), or *Elavl4* expression (D). Note especially the preferential *Elavl2* expression in a subset of GABAergic interneuron types.

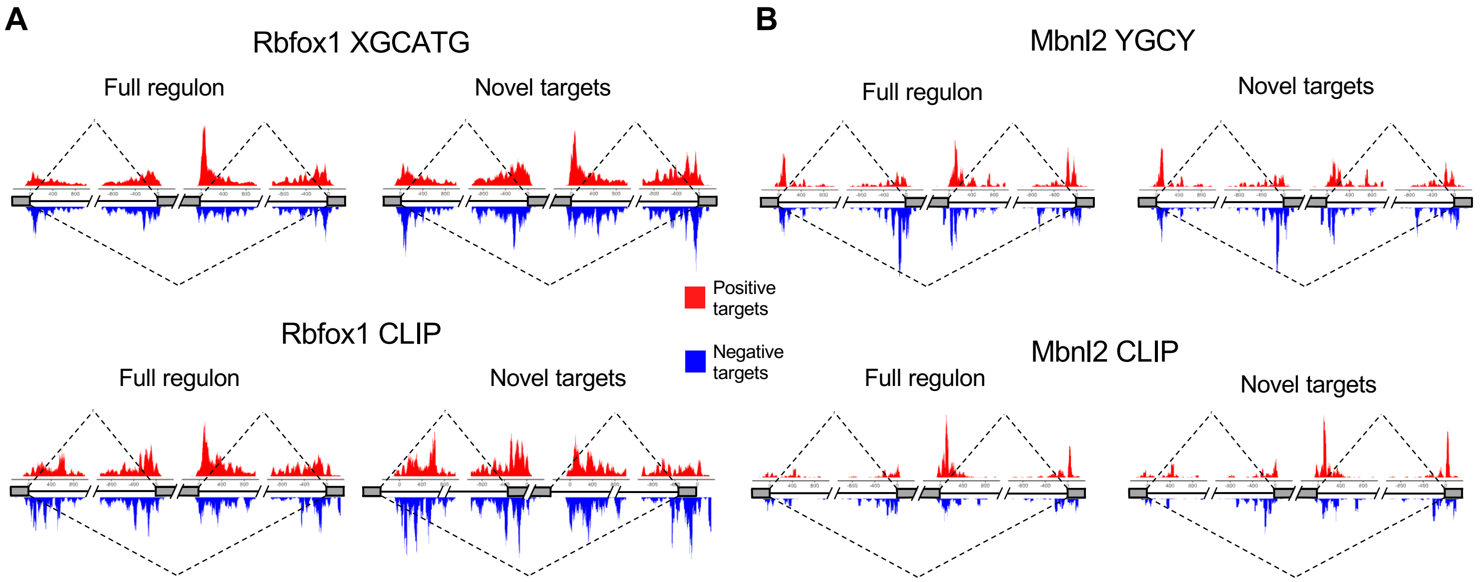


**Figure S3. RNAmap visualizations of CLIP tag and RBP binding sequence motif distributions. A,** Rbfox. **B,** Mbnl2. In each panel, normalized complexity RNA maps depict the distribution of RBP consensus binding motifs (top) and CLIP tags (bottom) at different positions relative to the alternative exons identified as positive targets (top, red) and negative targets (bottom, blue) of the RBP. Distributions are shown for both full regulons and novel targets alone (exons not previously identified by integrative modeling). Peaks just downstream of positively regulated exons and just upstream of negatively regulated exons are consistent with known position-dependent splicing regulation by these RBPs.

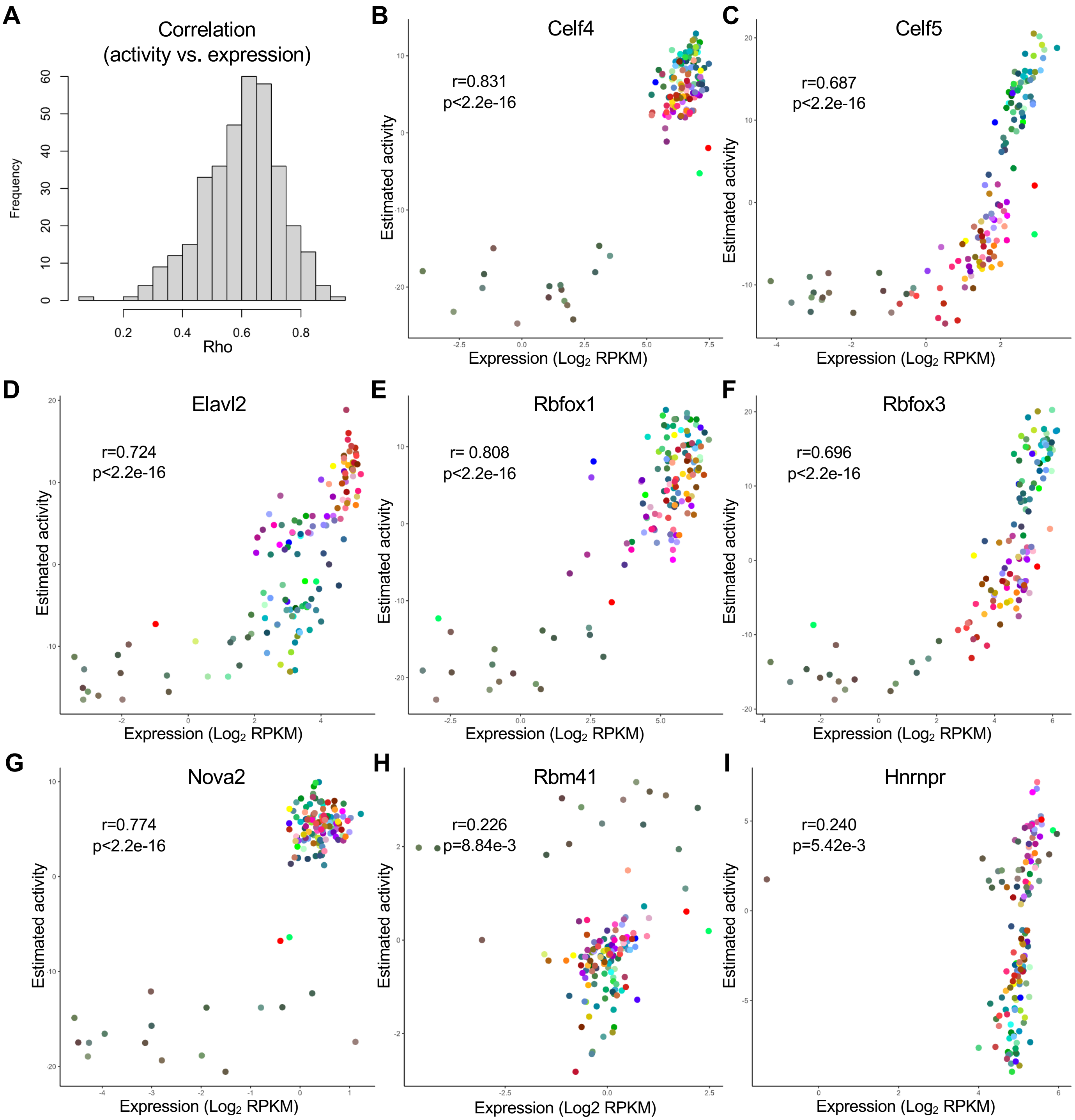


**Figure S4. Correlation between RBP activity and expression values. A**, Histogram showing the distribution of Spearman rank correlation coefficients ($\rho$). **B-I**, Representative RBPs with relatively high (B-G) or low (H-I) correlation.

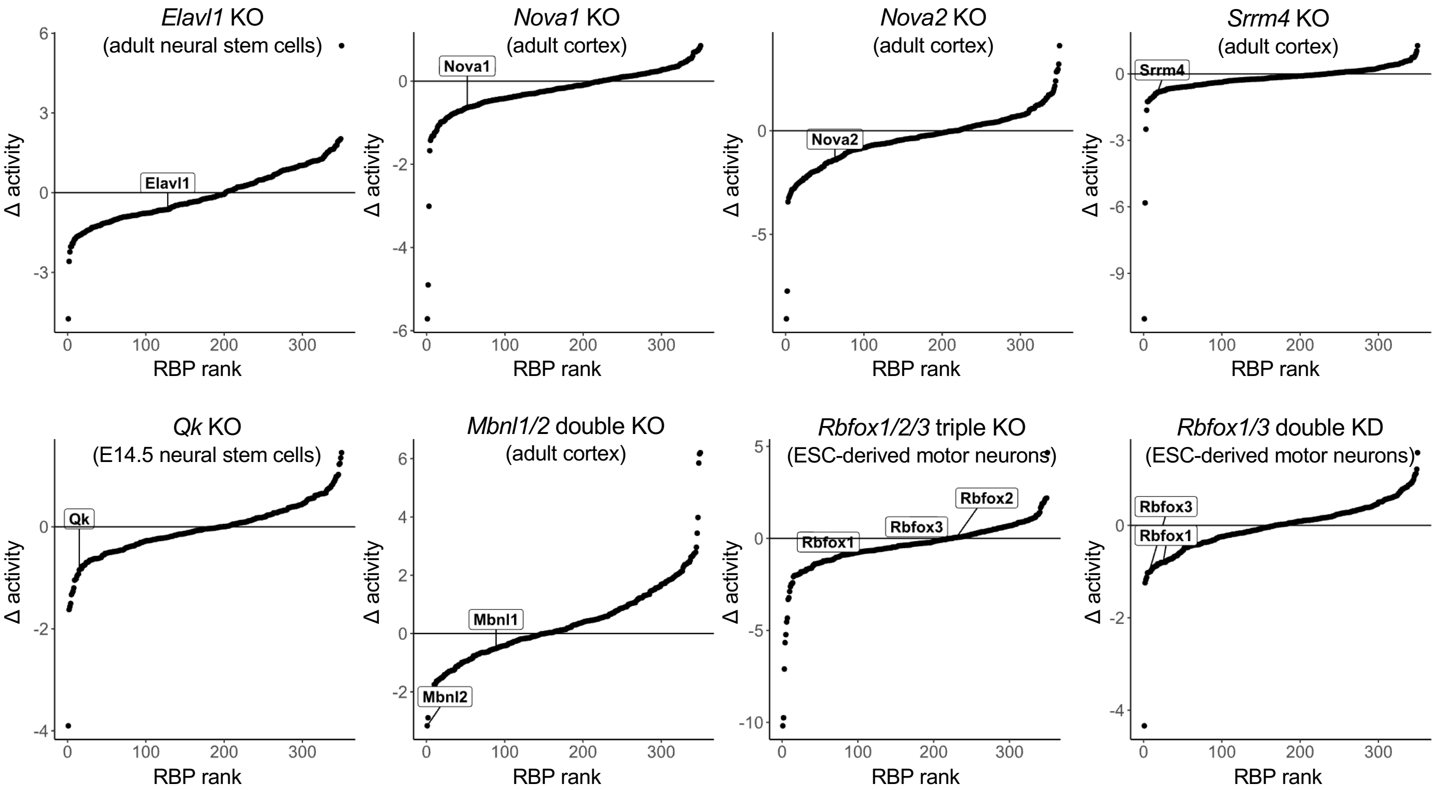


**Figure S5. Ranked differential RBP activity in RBP depleted vs. WT samples.** RBPs are ranked on the x-axis by activity difference (RBP depletion vs. WT) in various RBP knockout (KO) or RNAi knockdown (KD) data sets. Data sources: *Elavl1* KO: Wang et al. 2019; *Nova1* and *Nova2* KO: Saito et al. 2016; *Srrm4* KO: Quesnel-Vallieres et al. 2015; *Qk* KO: Hayakawa-Yano et al. 2017; *Mbnl1/2* double KO: Weyn-Vanhentenryck et al. 2018; *Rbfox1/2/3* triple KO: Jacko et al. 2018; *Rbfox1/3* double KD: Lee et al. 2016.

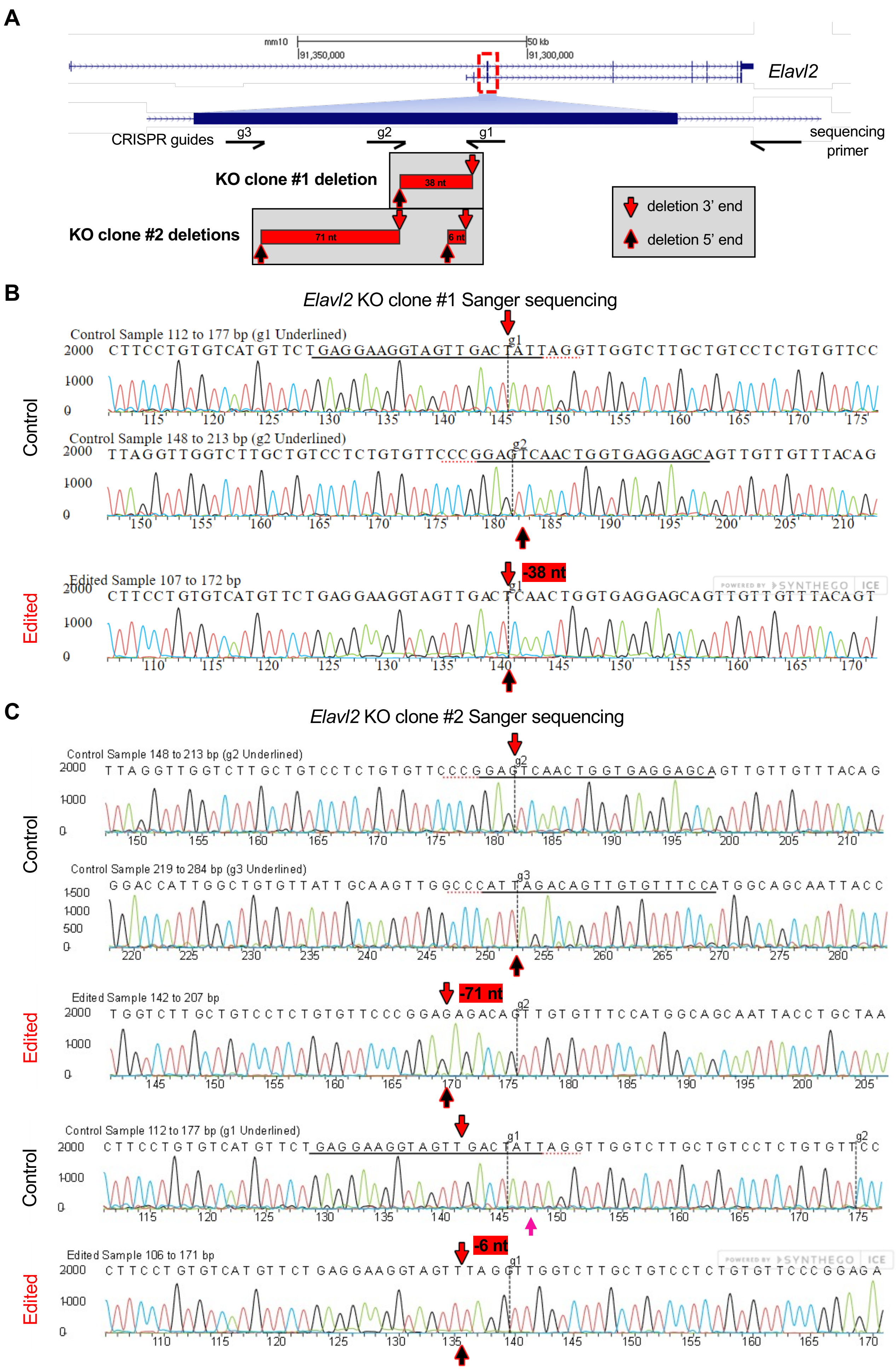

A
mm10
50 kb
91,350,000
91,300,000
Elavl2
CRISPR guides
g3
g2
g1
sequencing primer
KO clone #1 deletion
38 nt
KO clone #2 deletions
71 nt
6 nt
deletion 3' end
deletion 5' end
B
Elavl2 KO clone #1 Sanger sequencing
Control
Control Sample 112 to 177 bp (g1 Underlined)
Control Sample 148 to 213 bp (g2 Underlined)
Edited
-38 nt
Edited Sample 107 to 172 bp
C
Elavl2 KO clone #2 Sanger sequencing
Control
Control Sample 148 to 213 bp (g2 Underlined)
Control Sample 219 to 284 bp (g3 Underlined)
Edited
-71 nt
Edited Sample 142 to 207 bp
Control
Control Sample 112 to 177 bp (g1 Underlined)
Edited
-6 nt
Edited Sample 106 to 171 bp

**Figure S6. Sanger sequencing data for validation of *Elavl2* knockout. A**, CRISPR-Cas9 knockout design and resulting deletions. A set of three guides (from the Synthego CRISPR Gene Knockout Kit targeting *Elavl2*) were used to target the first constitutive exon of *Elavl2*, resulting in two clones with homozygous frameshifting deletions used in downstream experiments. The location of the downstream reverse-oriented primer used for Sanger sequencing is shown on the right. 5' and 3' deletion cut sites observed in two knockout clones are shown by pink and yellow arrows, respectively. **B**, Sanger sequencing of WT and *Elavl2* knockout clone 1 showing a 38 nucleotide (nt) frame-shifting deletion. The Synthego software ICE software was used for Sanger sequencing data visualization. **c**, Similar to (b) but for knockout clone #2, showing a 71 nt frame-shifting deletion as well as a 6 nt deletion.

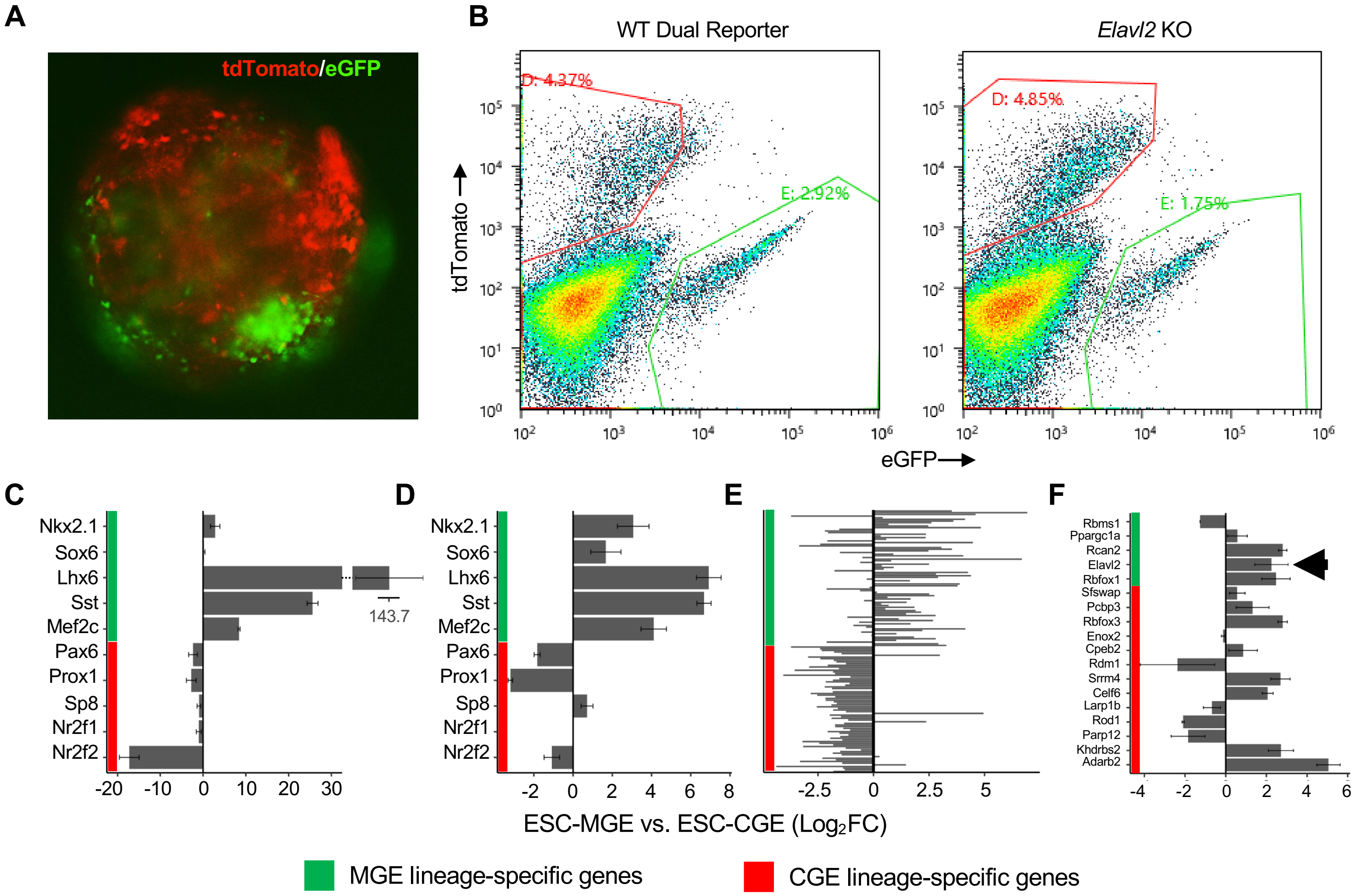


**Figure S7. Characterization of WT ESC-interneurons using MGE- and CGE-linage identity marker gene expression**. **A**, tdTomato and eGFP fluorescence in a day 12 EB. **B**, Representative scatter plots of FACS sort gating. **C,D**, qPCR (C) and RNAseq (D) of MGE- and CGE-specific canonical marker genes. **E**, Differentially expressed genes previously identified from comparison of MGE and CGE cells *in vivo* (Miyoshi et al. 2015) showed expected differences between ESC-MGE and ESC-CGE cells. **F**, Similar to (C-E) but for RBPs showing differential expression in adult MGE- vs CGE-lineage neurons in Tasic *et al.* 2018 data. Elavl2 expression difference is indicated by an arrow. Error bars represent standard error of the mean (SEM).

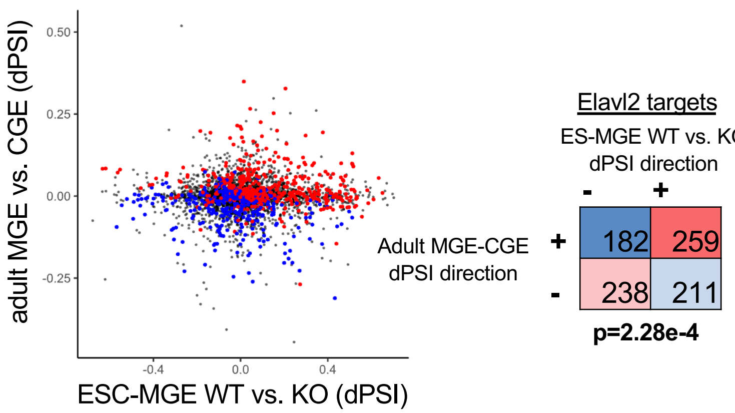


**Figure S8. Elavl2 knockout abrogates Elavl2 regulon activity and shifts ESC-MGE cells towards a CGE identity.** (Left) Exon inclusion differences (dPSI) between WT versus Elavl2 KO ESC-MGE cells on the x-axis and MGE versus CGE-lineage adult neurons on the y-axis. Elavl2 target exons are indicated by red or blue, respectively. (Right) Exact test of Elavl2 target exons shows the conchordance of directional splicing differences between the two comparisons, indicating the contribution of knockout-modulated exons to MGE identity.

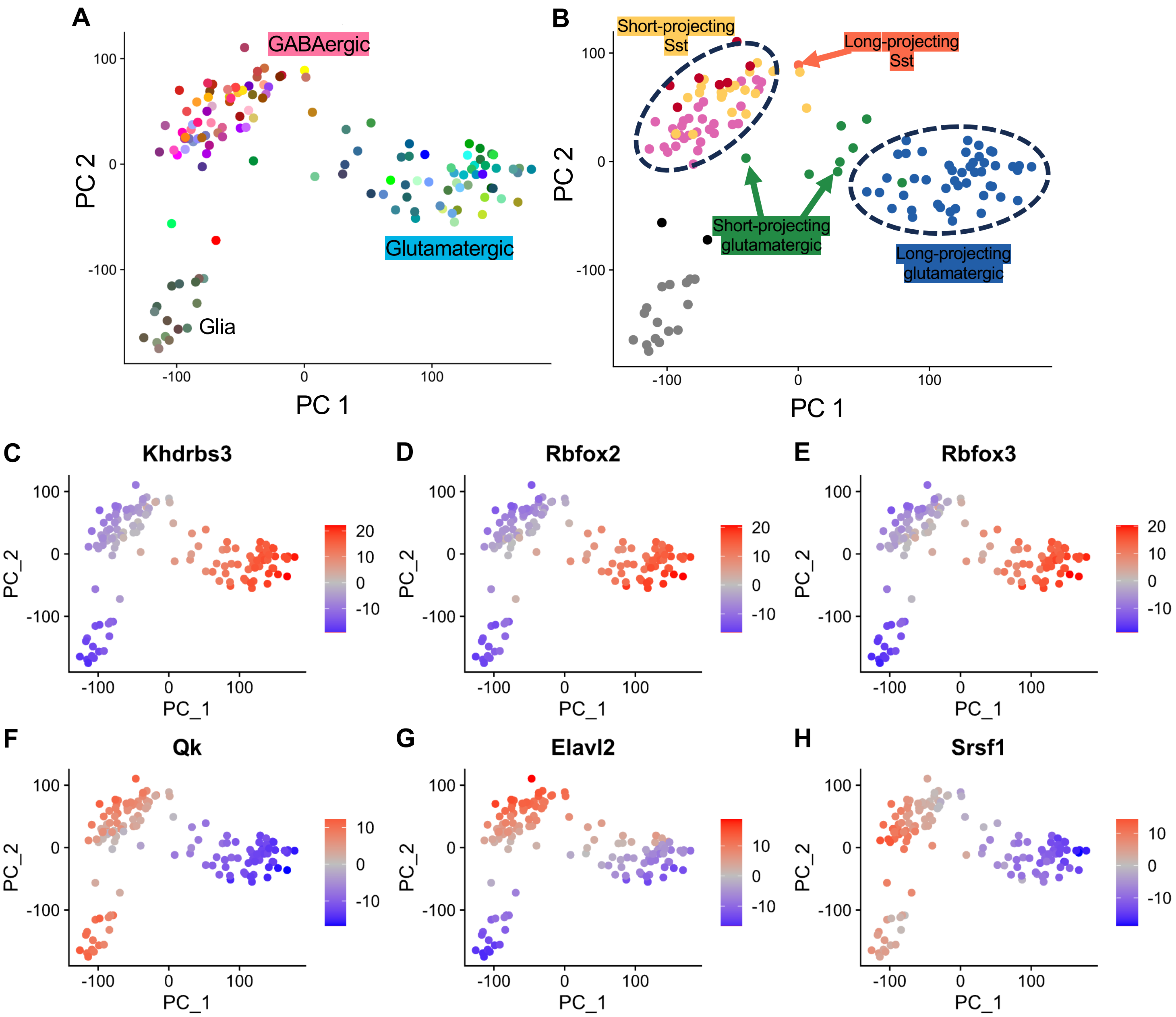


**Figure S9. PCA visualizations of RBP activity differences among neuron types. A**, PCA plot of the first two principal components of RBP activity with cell type clusters and neuronal classes indicated by color. **B**, Similar to (A) but with neuron projection types indicated by color. **C-H**, Similar to (A-B) but with neuron types colored by activity of long projecting neuron-specific (C-E) or short projecting neuron-specific activity (F-H).
